# A Synthetic Defective Interfering SARS-CoV-2

**DOI:** 10.1101/2020.11.22.393587

**Authors:** Shun Yao, Anoop Narayanan, Sydney Majowicz, Joyce Jose, Marco Archetti

## Abstract

Viruses thrive by exploiting the cells they infect but must also produce viral proteins to replicate and infect other cells. As a consequence, they are also susceptible to exploitation by defective versions of themselves that do not produce such proteins. A defective viral genome with deletions in protein-coding genes could still replicate in cells coinfected with full-length viruses, and even replicate faster due to its shorter size, interfering with the replication of the virus. We have created a synthetic defective interfering version of SARS-CoV-2, the virus causing the recent Covid-19 pandemic, assembling parts of the viral genome that do not code for any functional protein but enable it to be replicated and packaged. This synthetic defective genome replicates three times faster than SARS-CoV-2 in coinfected cells, and interferes with it, reducing the viral load of a cell by half in 24 hours. The synthetic genome is transmitted as efficiently as the full-length genome, confirming the location of the putative packaging signal of SARS-CoV-2. A version of such a synthetic construct could be used as a self-promoting antiviral therapy: by enabling replication of the synthetic genome, the virus promotes its own demise.

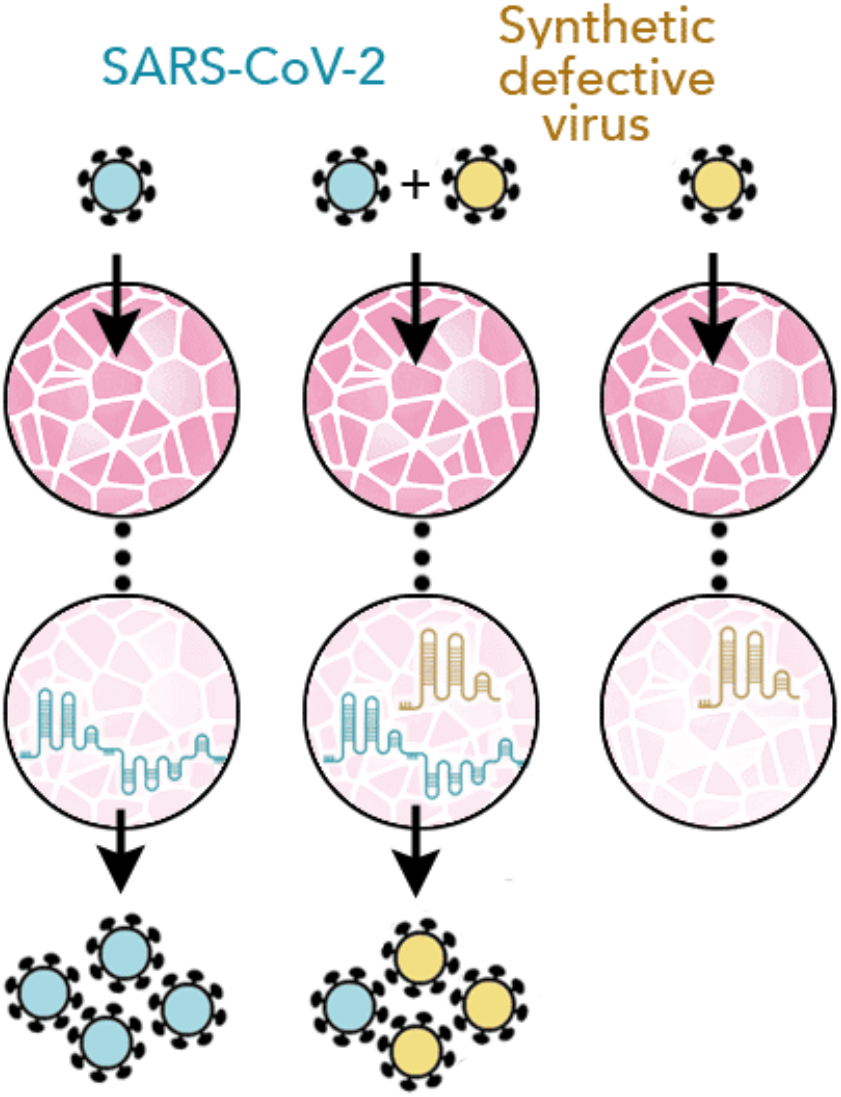
Graphic summary.

---

Versions of a viral genome with large deletions arise frequently from most RNA viruses when passaged in vitro [Gard et al. 1952, Brian & Spaan 1997, Vignuzzi & Huang & Baltimore 1970 López 2019]. Defective genomes lacking essential coding sequences can still replicate and be packaged into virions in the presence of functional full-length viruses that provide the essential proteins for replication and packaging, as long as the defective genome retains the ability to bind to these proteins (a similar process occurs when helper plasmids or bacteriophages enable replication or packaging of phagemids [Qi et al. 2012]). These defective genomes can be considered parasites of the full-length virus. Given their shorter length they have the potential to replicate faster than their parental full-length viral genome in coinfected cells and, because they compete for replication and packaging, to impair the growth and spread of the virus.

Such defective interfering (DI) genomes have been described – and indeed appear to be common – in coronaviruses [Kim et al. 1993a,b, Mendez et al. 1996, Brian & Spaan 1997, Izeta et al. 1999, Graham et al. 2005, Fehr & Perlman 2015, Vignuzzi & López 2019], where they have been used to locate the functional elements of their genomes, but not yet in SARS-CoV-2, the virus responsible for the current Covid-19 pandemic [Wu et al. 2020, Zou et al. 2020], even though deletions have been reported [Kim et al. 2020]. We made short synthetic DI genomes from parts of the SARS-CoV-2 genome, to test whether they could replicate in coinfected cells, be packaged into virions and impair the growth of the wild type (WT) virus.

The design of the DI genomes was based on observations from natural DI coronaviruses [TGEV: Mendez et al. 1996; MHV: Makino et al. 1985, 1988, 1990, Van der Most et al. 1991; Kim et al. 1993a, Kim et al. 1995, Masters 1994; Goebel et al. 2007; BCoV: Chang et al. 1994, Chang & Brian 1996; Williams et al. 1999; Raman et al. 2005; Brown et al. 2007; 229E: Thiel et al. 1998; IBV: Penzes et al. 1994, Dalton et al. 2001; reviewed by Yang & Leibowitz 2015] suggesting that the 3’ and 5’ untranslated regions (UTRs) are essential for replication and that the putative packaging signal resides inside the nsp15 sequence [TGEV: Escors et al. 2003, Morales et al. 2013; Hsieh et al. 2005; Hsin et al. 2018; MHV: Fosmire et al. 1992, Kuo & Masters 2013, Woo et al. 2019; BCoV: Cologna & Hogue 2000] – a conclusion that is however disputed for A betacoronaviruses, which lack the RNA structure responsible for packaging [Masters 2019].

Our main synthetic construct is made from three portions **[Figure 1]**: the 5’ UTR and the adjacent 5’ part of nsp1 in ORF1a; a part of nsp15 that includes the putative packaging signal; and the sequence spanning the 3’ part of the N sequence, ORF10 and the 3’UTR. We chose the N fragment to include two of the most conserved regions of the virus genome [Rangan et al. 2020] (28990..29054 and 28554..28569). Because there is evidence that a long ORF enables DIs in certain coronaviruses (notably MHV [De Groot et al. 1992], which is closely related to SARS-CoV-2) to replicate more efficiently (even if coding for a chimeric non-functional protein [Van der Most et al. 1995]), we assembled the three fragments in frame, to retain a 2247nt ORF starting at the 5’ end of nsp1 **[Figure 1]**; and because there is evidence [Joo & Makino 1995, VanMarle et al. 1995; Mendez et al. 1996] that multiple transcriptional regulatory sequences (TRS) reduce replication efficiency, we chose the 3’ portion to start from within the N sequence, to exclude its TRS. The synthetic sequence was analysed to check for potential, aberrations in the RNA secondary structure.

**Figure 1.**
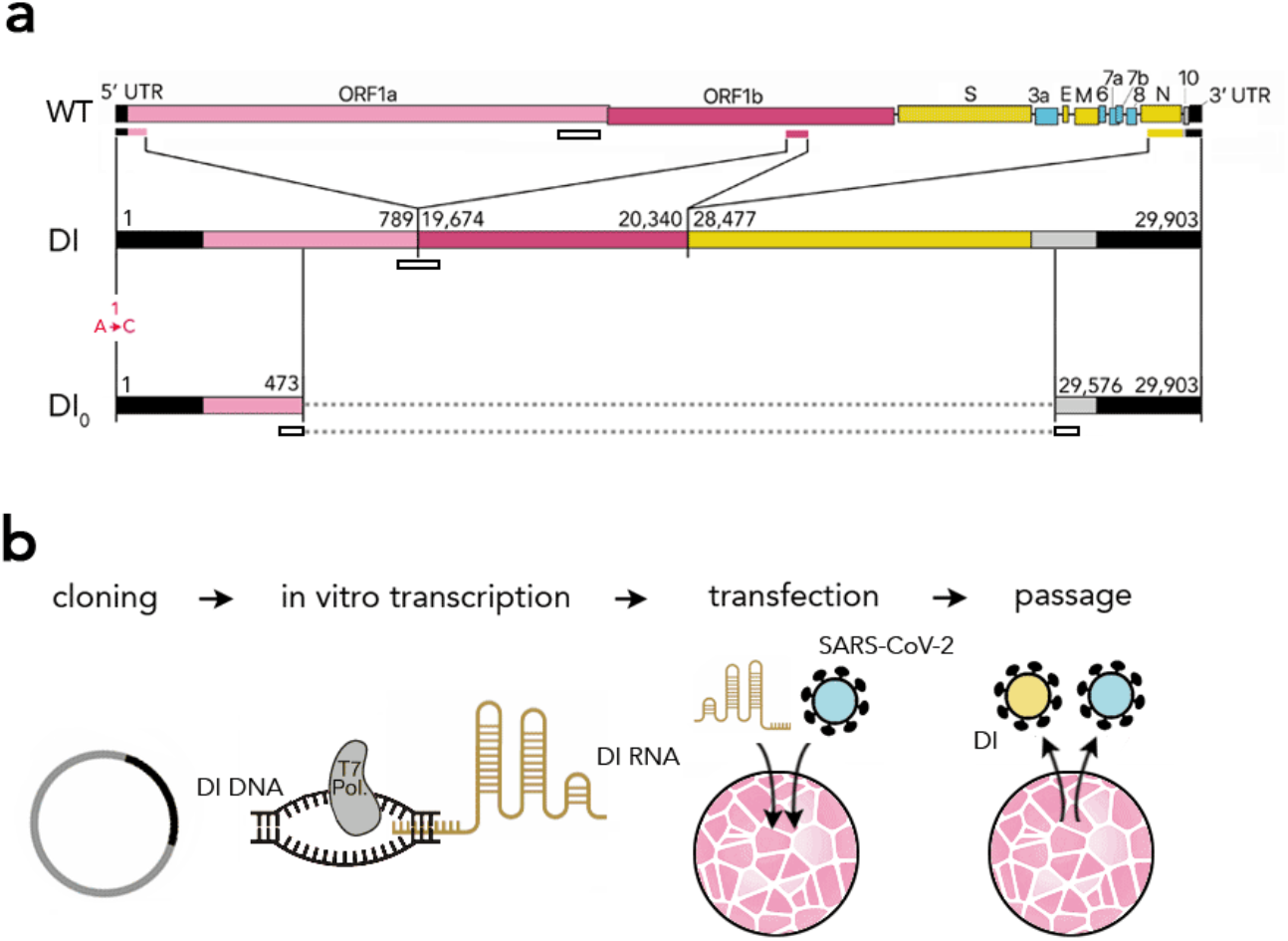
Synthetic defective interfering viruses. **a:** Three portions of the wild type (WT) SARS-CoV-2 genome were used to create a synthetic defective interfering genome (DI) and a shorter version (DI0) comprising only parts of the two terminal portions. Numbers delimiting the portions refer to positions in the SARS-CoV-2 genome. The first position is mutated (A→C) in both DI and DI_0_. Open rectangles show the position of the probes and primers used. **b**: In order to produce synthetic DI viruses, DNA constructs corresponding to the RNA sequence of DI or DI_0_ were transcribed into RNA in vitro using T7 RNA polymerase and transfected into Vero-E6 cells that were then infected with SARS-CoV-2. The supernatant from these cell cultures was used to infect new cells.

The length of our synthetic DI is 2882 nt, 9.6% of the full-length genome (29903 nt). We also synthetised a shorter (800 nt) defective genome (DI_0_) without the second portion (the putative packaging signal) and with shorter terminal portions **[Figure 1]**. The DI and DI_0_ genomes, synthetised as DNA and inserted into plasmids, were transcribed in vitro to yield genomic RNAs, which were then electroporated in Vero-E6 cells that were infected with SARS-CoV-2.

Because of the large amount of synthetic RNA transfected, the fast degradation of the synthetic RNA inside cells (in the absence of the virus, 1 to 4 % of the synthetic RNA can be detected by qRT-PCR 4 hours post transfection) and the lag between infection and the start of replication, it is not possible to quantify the replication rate of the DI and DI_0_ genomes, or even prove their replication, immediately after RNA transfection, as most of the RNA cannot replicate and will simply be degraded. It is possible, however, to quantify its effect on virus replication: the DI genome reduced the amount of SARS-CoV-2 by approximately half (compared to the amount of virus in control experiments) within 24 hours of transfection [**Figure 2a**]; the DI_0_ genome had no significant effect.

24 hours post transfection the supernatants were collected and used to infect new cell monolayers. In these cells we detected (by qRT-PCR) the DI and WT genomes, from 4 to 24 hours after the transfer. The DI_0_ genome was not detected, suggesting that the middle fragment of the DI synthetic sequence does have a positive effect on packaging. The transmission rate of the DI does not differ from that of the WT virus [**Figure 2b**], suggesting that the synthetic genome gets packaged into viral particles with essentially the same efficiency as the full-length virus, and that these viral particles are as infectious. In these cells coinfected by DI and SARS-CoV-2, the WT genome again declines by approximately half in 24 hours [**Figure 2c**]. The replication rate of the DI genome can now be quantified, revealing that it increases 3.3 times as fast as the WT virus (since the supernatant from the previous passage was removed 1 hours after infection, the increase we observed must be due to replication) [**Figure 2c**].

**Figure 2.**
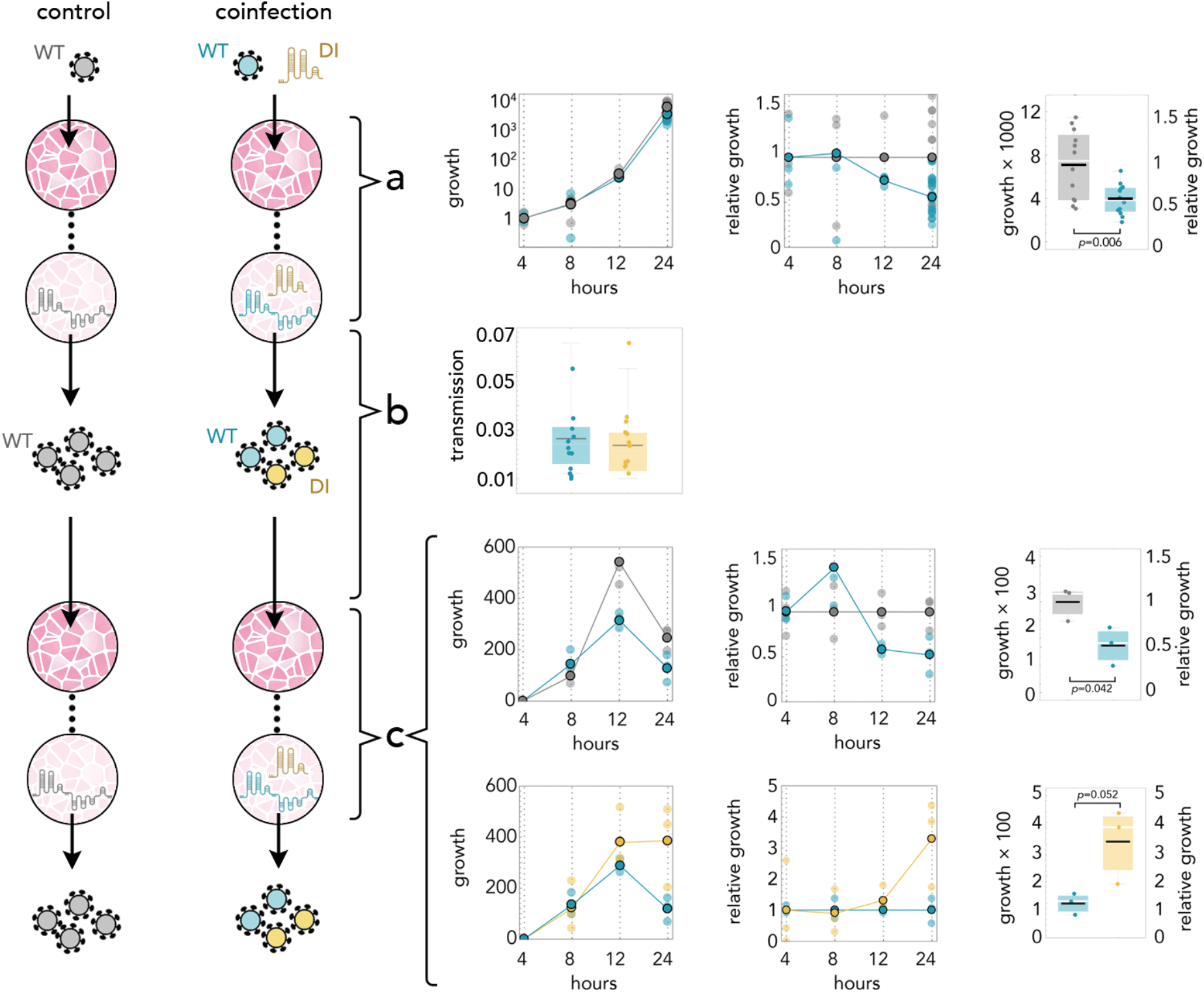
DI reduces the amount of SARS-CoV-2 by half; it replicates 3 times faster; and it is transmitted with the same efficiency. Yellow: DI in coinfections; blue: WT in coinfections; gray: WT in infections without DI. **a:** Growth rates (absolute amount relative to the amount at 4 hours) of WT in controls and in coinfections; growth relative to controls at the same time point; and detail at 24 hours. **b:** 24 hours after infection the supernatant was used to infect new cells. The transmission efficiency is the amount measured by qRT-PCR immediately before passaging divided by the average amount measured almost immediately (4 hours) after passaging. **c:** Growth rates (absolute amount relative to the amount at 4 hours) of WT in controls and in coinfections; growth relative to controls at the same time point; and detail at 24 hours. Growth rates (absolute amount relative to the amount at 4 hours) of WT and DI in coinfections; growth relative to that of WT in coinfections at the same time point; and detail at 24 hours.

We transferred the supernatant to new cells, coinfected with the WT virus, after 24 hours for four times. The DI genome was detected across all four passages, and the DI/WT ratio increased approximately 3 times every 24 hours, consistent with the relative replication advantage and equal transmission efficiency we measured. We were unable to measure the absolute WT/DI ratio because the amount of DI was below the level detectable by digital PCR. Our results, therefore, suggest that the interference of even a small amount of DI can have a strong impact on the replication of the WT virus and its burden for the cell.

To study the dynamics of the system we use a mathematical model of intra-cell competition between WT and DI genomes. Our model is inspired by a similar model [Shirogane et al. 2019] used to describe replication-independent poliovirus DIs; in our model, however, replication of the DI is entirely dependent on the WT. Such frequency-dependence can be thought of, in the framework of evolutionary game theory, as an interaction between cooperative (WT) and defector (DI) phenotypes engaged in a collective interaction for the production and exploitation of a public good (the viral proteins) [Szathmáry 1992, Turner & Chao 1999, Brown 2001]. Here we use the same approach but we take a step further by assuming that the replication of the DIs depends on the amount of proteins produced by multiple genomes (rather than two genomes engaged in a pairwise interaction [Szathmáry 1992, Turner & Chao 1999]) and that these have a nonlinear effect (rather than linear [Brown 2001]) on replication.

As predicted by early models of DI-WT dynamics [Szathmáry 1992] and by the general theory of non-liner public goods [Archetti 2018], a polymorphic equilibrium exists, in which DI and WT coexist, if the replication advantage (*R*) of the DI genome is below a critical threshold, whereas above that threshold DI drives WT to extinction. The critical threshold depends mainly on the number of genomes within the range of the viral protein produced by the WT genome (*n*) [**Figure 3**]. Other parameters, including the ones that control resource depletion, encapsidation and spontaneous decay, are less important for the dynamics (at the value of *R* we measured, for realistic values of *n,* they are almost irrelevant). While most of these parameters remain to be measured, therefore, the model suggests that the DI genome will, given the high replication advantage we measured, increase in frequency over time, eventually leading to the extinction of the WT genome [**Figure 3**]. As we monitored coinfections for only 5 passages, however, we were unable to verify whether the slight increase in DI/WT ratio we observed in coinfections is indeed the early stage of a reduction that would lead to the extinction of the WT virus.

**Figure 3.**
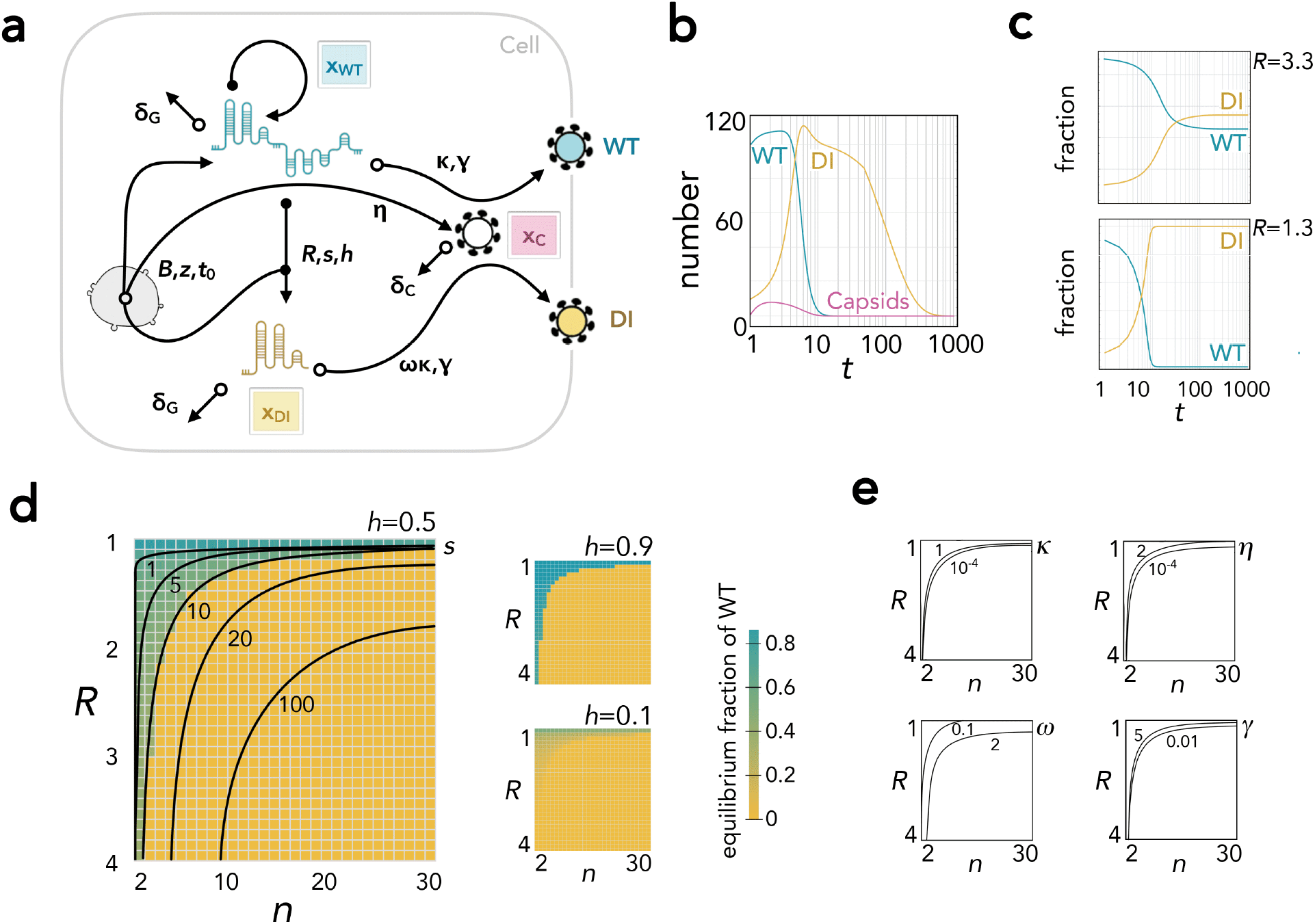
Simulation of the within-cell dynamics. **a: Flow diagram of the model**. The number of WT genomes (*x*_WT_), DI genomes (*x*_DI_) and capsids (*x*_C_) increase (full circles) due production and decline (empty circles) due to degradation and encapsidation. Production is proportional to the amount of resources of the cell (*B*), which decreases as a logistic function (with steepness *z* and inflection *t*_0_) of time (*t*); and increases as a linear function (for capsids) or as a logistic function (with steepness *s* and inflection *h*) of the number of WT genomes (for WT and DI genomes). DI genomes replicates at a rate *R* relative to WT genomes. Genomes decay at a rate δ_G_; capsids decay at a rate δ_C_. The rates of encapsidation are *κ* for WT genomes and *ωκ* for DI genomes; γ is the number of genomes per capsids; *η* is the capsid/genome ratio. **b: Example of the results: number of WT and DI genomes and capsids over time**. *R*=3.3 (the value measures empirically); *n*=10; *s*=10; *h*=0.5; *δ_G_*=0.01; *δ_C_*=0.001; *γ*=1; *η*=0.3; *κ*=0.03, ω=0.9; *B*=2, *z*=−0.03; *t*_0_=10; starting from 100 WT and 10 DI genomes. **c: Examples of the results: fraction of WT and DI genomes over time**. *R*=3.3 (the value measured empirically) or *R*=1.3, assuming no depletion of resources (*B*=2, *z*=0); *n*=10; *s*=10; *h*=0.5; *δ_G_*=0.01; *δ_C_*=0.001; *γ*=1; *η*=0.3; *κ*=0.03, ω=0.9. The initial fraction of DI in this example is 0.1, but the results are independent from the initial fraction. **d: Summary of the effect of***R* **and***n* **on the results.** The color of each cell shows the stable fraction of DI as a function of the replication advantage (*R*) of the DI genome and the number of genomes within the range of the viral protein produced by the WT genome (*n*), for different values of *h*; assuming no resource depletion (*B*=2, *z*=0); *s*=10; *δ_G_*=0.01; *δ_C_*=0.001; *γ*=1; *η*=0.3; *κ*=0.03, ω=0.9. For *h*=0.5 the curves show, for different values of s from 1 to 100, the separation between the combination of parameters for which WT goes extinct (below the curve) or remains at a stable polymorphic equilibrium (above). **e: Summary of the effect of other parameters on the results**. Combinations of *R* and *n* below the curves lead to the extinction of WT. The curves are drawn for different values of *γ*, *η*, *κ* and ω; other parameters now shown because differences are undetectable. In all cases, *h*=0.5, *s*=10.

DI particles have long been known to virologists [Gard et al. 1952, Huang & Baltimore 1970] and their use in unravelling the location of functional elements of a genome is well known. Our synthetic DIs reveals that a disputed [Masters 2019] putative packaging sequence of SARS-CoV-2 does indeed enable packaging of our synthetic defective genome – and therefore presumably acts as a packaging signal for the WT genome. However, because the difference between our DI and DI_0_ synthetic constructs is not limited to the portion with the putative packaging signal, we cannot rule out that the packaging signals resides in the other parts of DI that DI_0_ lacks, most notably a conserved region (28554..28569) with a SL5 motifs in the N partial sequence included in the DI genome but not in the DI_0_ genome.

The interference with the WT virus is the most remarkable effect of our DI construct. DI particles are often described as by-products of inefficient replication or as having a regulatory function. Seen instead as defectors in the sense of evolutionary game theory [Szathmáry 1992, Turner & Cao 1999, Brown 2001], they need not serve any purpose for the WT virus: they exist as ultra-selfish replicators, able to free-ride as parasites of the full-length genome. As such, DI particles have a potential as antivirals: by virtue of their faster replication in cells coinfected with the virus, DI genomes can replicate faster and, in the process, interfere with the WT virus. As the DI genomes increase in frequency among the virus particles pool, the process becomes more and more effective, until the reduction in the amount of WT virus leads to the demise of both virus and DI. By enabling the replication and spread of the DI genome, the virus promotes its own demise. The potential of DIs as antivirals has been suggested, but not yet exploited, for other viruses [Marriott & Dimmock 2010, Dimmock & Easton 2014, Vignuzzi & López 2019], and a similar therapeutic approach has been proposed for bacteria [Brown et al. 2009] and cancer [Archetti 2013, Archetti et al. 2015]. While viruses such as HIV and influenza where DI therapy has been attempted [Marriott & Dimmock 2010, Dimmock & Easton 2014, Vignuzzi & López 2019] are not ideal for this approach, because of their short genome, multiple genomic fragments and complex life cycles, coronaviruses are ideal candidates because of their long single strand RNA genome and relatively simple replication process. A version of our synthetic DI could be used as an antiviral that would be self-sustaining and evolution-proof, or as a self-disseminating approach to suppress zoonoses [Nuismer & Bull 2020] that could spill over to humans in the future.

## Methods

### Sequences and cloning

The DNA sequence of the DI genome (GenBank accession number: MW250351) was designed to correspond to the following three joint portion s of the SARS-CoV-2 complete genome (the NCBI Reference Sequence for SARS-CoV-2; GenBank accession number: NC_045512.2), in the following order: 1 to 789; 19674 to 20340; and 28477 to 29903. The DI_0_ genome (GenBank accession number: MW250350) was designed to correspond to the following two joint fragments of SARS-CoV-2 in the following order: 1 to 473; 29576 to 29903. In both cases, the first nucleotide of the first fragment was changed from A to C in order to improve in vitro transcription efficiency [Milligan 1987, Martin & Coleman 1987]. The synthetic sequence was analysed using the Vienna RNA package [Lorenz et al. 2011] to confirm the absence of potential aberrations in the RNA secondary structure. The DI and DI_0_ genome DNA were assembled from synthetic oligonucleotides and inserted into a pMA-RQ plasmid by Invitrogen (Thermo Fisher Scientific). The T7 promoter TAATACGACTCACTATAGG was synthetised immediately upstream of the 5’ end of the synthetic virus sequence. A short sequence (CCATGG) containing the NcoI restriction site was synthetised immediately upstream of the 5’ end of the T7 promoter, and a short sequence (CCGGT) containing the AgeI restriction site was synthetised immediately downstream of the 3’ end of the third fragment. The plasmid DNA was purified from transformed bacteria and the final construct was verified by sequencing.

### In vitro transcription

The plasmid containing the synthetic DI or DI_0_ genomed DNA was linearized using NcoI and AgeI and resuspended in H_2_O. 1 μg was then used as a template to produce capped RNA via T7 RNA polymerase, using a single reaction setup of the mMESSAGE mMACHINE® Kit (Applied Biosystems), which contains: 2 μL enzyme mix (buffered 50% glycerol containing RNA polymerase, RNase inhibitor, and other components); 2 μL reaction buffer (salts, buffer, dithiothreitol, and other ingredients); 10 μL of a neutralized buffered solution containing: 15 mM ATP, 15 mM CTP, 15 mM UTP, 3mM GTP and 12mM cap analog [m7G(5’)ppp(5’)G]; 4 μL nuclease-free H_2_O; incubated for 2 hours at 37°C. RNA was isolated using TRIzol reagent (Invitrogen) extraction and isopropanol precipitation.

### Cells and transfection

Vero-E6 cells (*Chlorocebus sabaeus* kidney epithelial cells [Yasumura & Kawakita 1963]) cultured in DMEM medium (Hyclone, #SH30022.FS) supplemented with 10% fetal bovine serum (Corning, #35-011-CV), 100 units ml^−1^ penicillin and 100 μg ml^−1^ streptomycin (Gibco, #15140122) maintained at 37 °C and in a 5% CO_2_ atmosphere were grown to 80% confluence. The cells were electroporated with the RNA produced by in vitro transcription (DI: 532ng; DI_0_: 476ng; per 200,000 cells; equivalent to 1.7×10^6^ and 5.6×10^6^ RNA molecules per cell, respectively), in 100 μl Nucleocuvette Vessels using the SF Cell solution and program DN-100 on a 4D Nucleofector X unit (Lonza). The efficiency of transfection was approximately 90%. Cells used for the control experiments were electroporated in the same way but without RNA.

### Virus culture

SARS-CoV-2 isolate USA-WA1/2020 was obtained from BEI resources (#NR-52281) and propagated in Vero-E6 cells. Virus stocks were prepared and the titer as determined by plaque assays by serially diluting virus stock on Vero-E6 monolayers in the wells of a 24-well plate (Greiner bio-one, #662160). The plates were incubated at room temperature in a laminar flow hood with hand rocking every ten minutes. After one hour, an overlay medium containing 1XMEM, 1% Cellulose (Millipore Sigma, #435244), 2% FBS and 10mM Hepes 7.5 was added and the plates were incubated for a further 48 hours at 37^0^C. The plaques were visualized by standard crystal violet staining. All work with the SARS-CoV-2 was conducted in Biosafety Level-3 conditions at the Eva J Pell Laboratory of Advanced Biological Research, The Pennsylvania State University, following the guidelines approved by the Institutional Biosafety Committee.

### Coinfection and RNA extraction

200,000 transfected cells were seeded in each well of a 24-well plate (each well in triplicate), and incubated for 1 hour before being inoculated with SARS-CoV-2 at MOI=10. The medium containing the infectious SARS-CoV-2 viruses was removed after 1 hour and replaced with fresh medium. Cells were allowed to grow for 4, 8, 12 or 24 hours before RNA was extracted. The supernatant of cultures grown for 24 hours was used to infect new cells in 24-well plates for one hour, then media was replaced with fresh media and RNA was extracted from the cells after another 24 hours. This step was repeated four times to obtain RNA from four consecutive passages. RNA was extracted using Quick RNA miniprep kit (Zymo, #R1055) or TRIzol reagent (Invitrogen, #15596026) followed by isopropanol precipitation.

### RNA analysis

Equal amounts of total RNA were reverse transcribed into first-strand cDNA using Revert Aid™ First Strand cDNA Synthesis Kit (Fermentas). 2 μl diluted cDNA (3pg-100ng depending on the experiment) mixed 2 μl of 5 μM primer mix (forward plus reverse), 1 μl of 2 μM of probe and 5 μl master mix (2×) was used for qRT-PCR using TaqMan assay on a StepOnePlus instrument (Applied Biosystems) starting with polymerase activation at 95°C for 3 minutes, followed by 40 cycles of denaturation (95°C, 15 seconds) and annealing/extension (60°C, 1 minute). The amount of WT and synthetic DI or DI_0_ genome were quantified (using StepOnePlus Software 2.3) by the comparative C_T_ method [Livak and Schmittgen 2001]. All results were normalised with reference to the actin beta (ACTB) gene of *Chlorocebus sabaeus*; each sample was repeated three times and the average value was used; all absolute values reported are 2^−ΔCT^ values. Primers and probes for the DI and DI_0_ genomes were designed to amplify one of the junctions between portions of the WT genome [**Figure 1**]; for the virus we used a modified version of the CCDC primer-probe set on ORF1. A BLAST search revealed no off-target sequences neither in the SARS-CoV-2 nor in the *Chlorocebus sabaeus* genome. Primers and probes for the DI and DI_0_ genomes, for the SARS-CoV-2 genome, and for the ACTB gene of Vero-E6 cells, were labelled using the FAM dye, an IBFQ quencher and an additional internal (ZEN) quencher, and were synthetised by Integrated DNA Technologies. The sequences are the following:

DI:

Forward: 5’-AGCTTGGCACTGATCCTTATG-3’
Reverse: 5’-ACATCAACACCATCAACTTTTGTG-3’
Probe: 5’-FAM/TTACCCGTGAACTCATGCGACAGG/IBFQ-3’
DI_0_:

Forward: 5’- ATCAGAGGCACGTCAACATC –3’
Reverse: 5’- TTCATTCTGCACAAGAGTAGACT –3’
Probe: 5’-FAM/ AGCCCTATGTGTCGCTTTTCCGT /IBFQ-3’
SARS-CoV-2:

Forward: 5’- CCCTGTGGGTTTTACACTTAA –3’
Reverse: 5’- ACGATTGTGCATCAGCTGA –3’
Probe: 5’-FAM/CCGTCTGCGGTATGTGGAAAGGTTATG /IBFQ-3’
ACTB:

Forward: 5’-AGGATTCATATGTGGGCGATG-3’
Reverse: 5’-AGCTCATTGTAGAAGGTGTGG-3’
Probe: 5’-FAM/AGCACGGCATCGTCACCAACT/IBFQ-3’

### Mathematical model

We model the dynamics of intracellular competition between DI and WT using the following ordinary differential equation system:

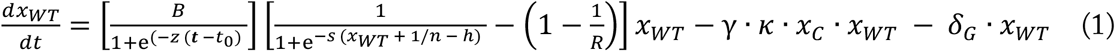

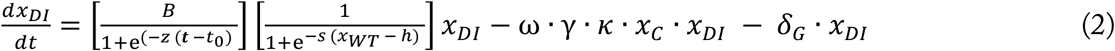

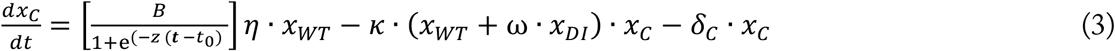

The three equations (1–3) represent the change over time (*t*) of, respectively, the number of WT genomes (*x*_WT_), of DI genomes (*x*_DI_) and capsids (*x*_C_). In each equation, the first term is the rate of increase due to replication; the second is the rate of loss due to encapsidation; the third is the rate of loss due to degradation. New WT and DI genomes are produced (first term of equations 1 and 2, respectively) at a rate given by a logistic function (with steepness *s* and inflection point *h*) of the number of WT genomes (as DI genomes do not produce any viral protein) multiplied by a factor corresponding to the amount of resource (*B*) within the cell, which we assume is time-dependent and changes at a rate given by a logistic function with negative steepness *z* and inflection point *t_0_* (the time point when resources are depleted by half). WT genomes pay a cost equal to 1-1/*R*, where *R* is the ratio between the rate of replication of DI and WT (*R*>1 given that the DI genome is shorter and can replicate faster) but have a slight advantage due to the additional viral genome (itself) producing replication proteins among the *n* neighbouring genomes. We assume that *n* is large enough that we can ignore the variance in the number of WT genomes around *n* (which, in a large population, would be binomially distributed). WT and DI genomes decrease as a function of their decay rate *δ_G_* and the rate of encapsidation (*κ*, the rate of encapsidation for the WT genome; ω, the ratio between the rate of encapsidation of DI and WT; and *γ*, number of genomes per capsids). The number of capsids (equation 3) increase as a linear function (controlled by the capsid/genome ratio *η*) of the number of WT genomes and decreases as a function of the number of encapsidated genomes; and as a function of the decay rate *δ_C_*.

## Contributions

Project design and coordination: MA

Sequence design: MA

Cloning: SY, MA

In vitro transcription: SY, MA

Transfection: SY, MA

Virus culture and infection: JJ, AN, SM

RNA extraction: AN, SM, SY

qRT-PCR: SY, MA

Data analysis: MA, SY

Mathematical model: MA

Writing: MA

## Conflict of interests

Pending patent application (MA and Pennsylvania State University).

## Funding

MA, JJ startup; Huck COVID-19 seed grant.

